# Mapping individualized multi-scale hierarchical brain functional networks from fMRI by self-supervised deep learning

**DOI:** 10.1101/2025.04.07.647618

**Authors:** Hongming Li, Chuanjun Zhuo, Zaixu Cui, Matthew Cieslak, Taylor Salo, Raquel E. Gur, Ruben C. Gur, Russell T. Shinohara, Desmond J. Oathes, Christos Davatzikos, Theodore D. Satterthwaite, Yong Fan

## Abstract

The brain’s multi-scale hierarchical organization supports functional segregation and integration. Characterizing the hierarchy of individualized multi-scale functional networks (FNs) is crucial for understanding these fundamental brain processes. It provides promising opportunities for both basic neuroscience and translational research in neuropsychiatric illness. However, current methods typically compute individualized FNs at a single scale and are not equipped to quantify any possible hierarchical organization. To address this limitation, we present a self-supervised deep learning (DL) framework that simultaneously computes multi-scale FNs and characterizes their across-scale hierarchical structure at the individual level. Our method learns intrinsic representations of fMRI data in a low-dimensional latent space to effectively encode multi-scale FNs and their hierarchical structure by optimizing functional homogeneity of FNs across scales jointly in an end-to-end learning manner. A DL model trained on fMRI scans from the Human Connectome Project successfully identified individualized multi-scale hierarchical FNs for unseen individuals and generalized to two external cohorts. Furthermore, the individualized hierarchical structure of FNs was significantly associated with biological phenotypes, including sex, brain development, and brain health. Our framework provides an effective method to compute multi-scale FNs and to characterize the inter-scale hierarchy of FNs for individuals, facilitating a comprehensive understanding of brain functional organization and its inter-individual variation.

## Introduction

Functional magnetic resonance imaging (fMRI) is a powerful tool for studying the organization of functional networks (FNs) in the human brain. Recent work has demonstrated that these FNs are unique to each individual [1-10], creating unprecedented translational opportunities when combined with large-scale neuroimaging datasets. This individualized mapping of FNs has the potential to revolutionize the development of new diagnostics for neuropsychiatric illnesses, ultimately leading to neuromodulatory therapies targeted according to each individual’s unique functional neuroanatomy. This potential is further strengthened by recent evidence that inter-individual variation in FNs is linked to development in youth [3], sex differences [11], individual differences in cognitive performance [12], and trans-diagnostic psychopathology [13].

A variety of techniques are available for computing individualized FNs [6, 14-26]. These methods typically employ explicit constraints or regularizations to maintain the spatial correspondence of FNs across individuals. Among them, independent component analysis (ICA)-based methods [24-26] promote similarity between individualized and group level FNs. Other methods utilize constraints based on assumed statistical distributions of FN loadings across individuals [6, 18, 21, 23]. Spatially-regularized non-negative matrix factorization (NMF) relaxes assumptions about these statistical distributions. More recently, we developed a self-supervised deep learning (DL) method to compute individualized FNs while implicitly maintaining inter-individual correspondence through predictive modeling [27].

Despite these promising findings on individualized FNs [1-10], a key limitation of current methods is their reliance on the computation of FNs at a single scale, which is ineffective for characterizing the brain’s inherent multi-scale hierarchical functional organization [28-36]. The brain functional hierarchy facilitates the segregation and integration of brain functional networks across different spatial scales, enabling efficient information processing and supporting complex function [28, 34]. Prior work has shown that associations of FN topography with development and cognition may vary by scale [37]. Therefore, characterizing these multi-scale hierarchical FNs at the individual level could provide insights into inter-individual variation in this hierarchy and its potential links to biological, clinical, and behavioral phenotypes.

Clustering and module detection algorithms can help detect the hierarchical organization of FNs [30-34]. However, their performance depends on pre-defined network nodes or FNs. Deep learning methods like autoencoders show promise for deriving multi-scale, low-dimensional representations of FNs [38-42], but have primarily been used at the group level. A recently developed telescopic ICA strategy builds a hierarchy of multi-scale FNs by recursively applying ICA [43], but this method only captures a group-level hierarchy and is prone to error accumulation due to its recursive nature.

To address these limitations, we developed a self-supervised DL framework to compute multi-scale FNs and quantify their hierarchical organization at the individual level. We hypothesized that learning a low-dimensional latent representation of fMRI data would facilitate computing fine-scale individualized FNs and inferring inter-scale hierarchies, which are quantified by hierarchy coefficients (HCs) for characterizing the integration of FNs from fine to coarse scales and enabling the identification of multi-scale FNs with a hierarchical structure. We used convolutional neural networks (CNNs) with an encoder-decoder architecture to jointly compute fine- and coarse-scale FNs and infer their hierarchy by aggregating fine-scale FNs to form coarse-scale FNs. This framework trains a DL model by optimizing a self-supervised loss function that measures the functional homogeneity of FNs at all scales jointly. The trained model can then compute individualized, multi-scale hierarchical FNs in a single forward pass on new fMRI data. Leveraging resting-state fMRI scans from the Human Connectome Project (HCP) [44], we trained a model to compute FNs across a three-scale hierarchy, comprising 148, 50, and 17 FNs at fine, intermediate, and coarse scales, respectively. This model enabled the computation of individualized multi-scale FNs and the characterization of their hierarchical structure within individuals. Results demonstrated that the identified FNs aligned with known large-scale FNs at the coarse scale and the model generalized well to external datasets. Furthermore, the individualized hierarchies of FNs were significantly associated with biological phenotypes, including sex, brain development, and schizophrenia diagnosis, highlighting their potential as quantitative measures of individualized brain functional network organization. This work provides an effective approach to compute multi-scale FNs and quantitatively characterize their inter-scale hierarchies at the individual level, facilitating a comprehensive understanding of brain functional organization and its inter-individual variation.

## Results

The proposed DL framework consists of three modules for representation learning, FN learning, and functional hierarchy learning (Fig. 1a). The representation learning module learns time-invariant feature maps from the input fMRI data, accounting for the temporal misalignment of resting-state fMRI scans from different sessions and individuals, as used in [27] (Fig. 1b). The FN learning module is a network with an encoder-decoder architecture: the encoder learns a low-dimensional latent embedding of the fMRI data, informative for computing the multi-scale FNs and their hierarchical structure; and the decoder identifies the FNs at the finest scale (Fig. 1c). The functional hierarchy learning module is a network to infer the HCs between consecutive scales from the latent embeddings obtained in the FN learning module (Fig. 1d). The HCs quantify the hierarchical structure of FNs between scales, according to which finer-scale FNs are aggregated into coarser-scale FNs (Fig. 1a). The DL model is trained with self-supervision measures, including functional homogeneity and regularization at all scales, so that the multi-scale hierarchical FNs are optimized jointly without requiring any external supervision. Details about network architecture and model implementation are presented in Methods.

**Fig. 1.**
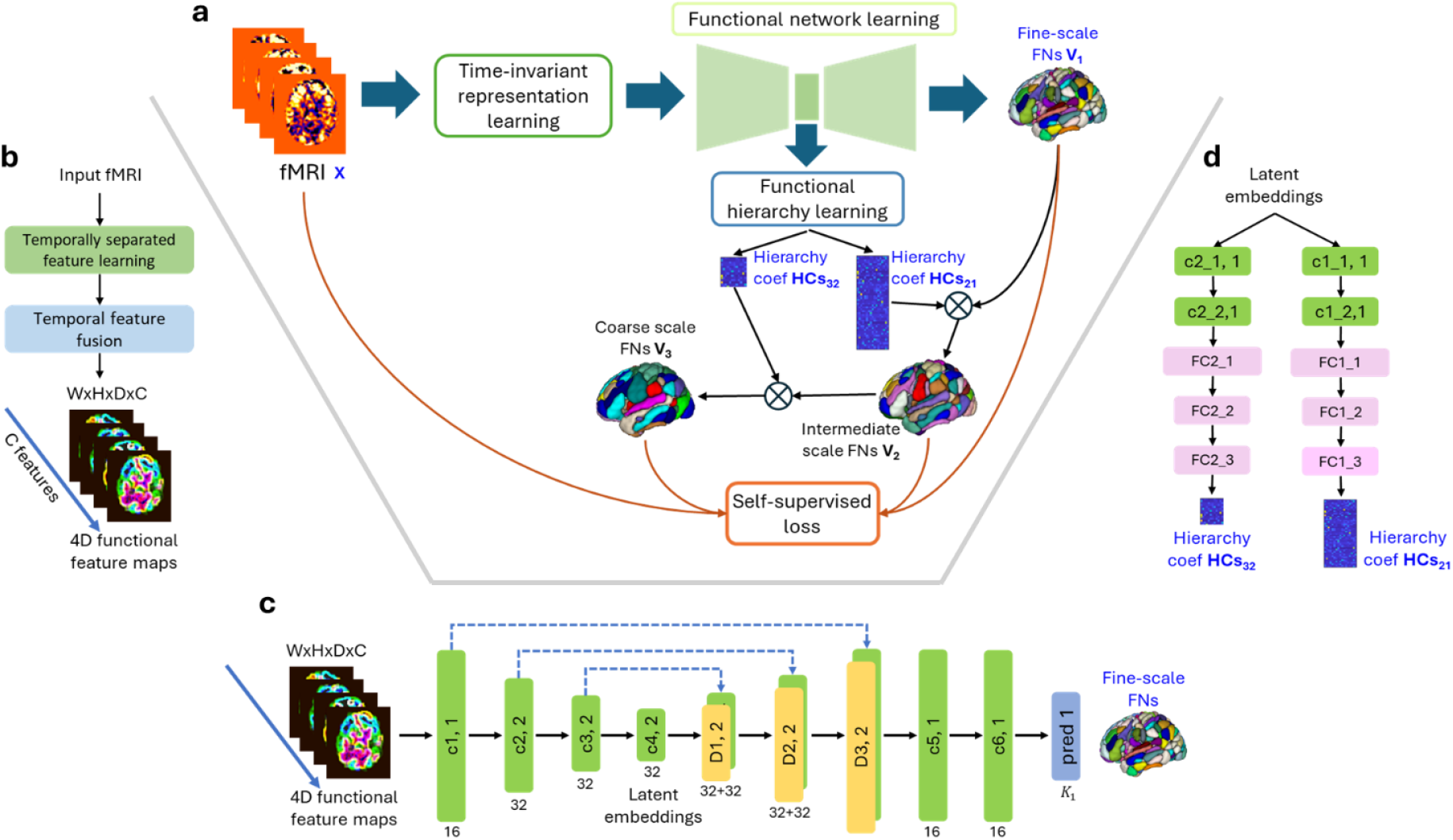
DL model for computing individualized, multi-scale hierarchical FNs. (**a**) Schematic diagram of the DL model for computing a three-scale hierarchy with *K*_1_, *K*_2_, and *K*_3_ FNs at fine, intermediate, and coarse scales, respectively. It consists of a time-invariant representation learning module, a functional network learning module, and a functional hierarchy learning module. (**b**) The time-invariant representation learning module learns informative features to account for temporal misalignment of resting-state fMRI data across different scans. Different colormaps are used to distinguish the input fMRI data from the feature maps learned by the module; the colors are for illustrative purposes only. (**c**) Network architecture of the functional network learning module for computing individualized FNs at the fine scale. The numbers underneath convolutional (c1 to c6, green blocks) and deconvolutional (D1 to D3, yellow blocks) layers indicate their corresponding numbers of kernels with a stride of 1 or 2. (**d**) Network architecture of the functional hierarchy learning module to learn the HCs between two consecutive scales (from fine to intermediate, and from intermediate to coarse). The hierarchy learning module consists of two branches, each with two convolutional layers and three fully connected (FC) layers, to infer the FN relationships between consecutive scales. Across scales, the finer-scale FNs are aggregated jointly into coarser-scale FNs according to the HCs.

### Self-supervised DL model could identify individualized hierarchical FNs

We trained a DL model using resting-sate fMRI data of 678 individuals randomly selected from the HCP cohort. The numbers of FNs at fine, intermediate, and coarse scales were set to 148, 50, and 17, respectively, based on a data-driven estimation and the popular settings in large-scale FN modeling. The trained DL model was then applied to fMRI data from an independent set of 400 HCP individuals for evaluation.

Fig. 2 illustrates functional parcellations derived from the group average FNs of the 400 testing individuals, along with those from three randomly selected individuals, across the fine, intermediate, and coarse scales. The FNs exhibit a nested hierarchical structure from fine to coarse scales. For example, the fine-grained regions visible at the finest scale within the sensorimotor network were aggregated into larger regions at the intermediate, and coarser scales. The identified FNs at the coarse scale included both spatially localized FNs, such as visual and somatomotor networks, and distributed FNs, such as default mode and frontoparietal networks. The coarse-scale FNs identified by the DL model showed statistically significant alignment with the canonical networks from the Yeo atlas [45] (*p* < 0.001), as determined by a spin-based spatial permutation procedure [46]. All group average FNs from fine to coarse scales are illustrated in Figs. S1-S3.

**Fig. 2.**
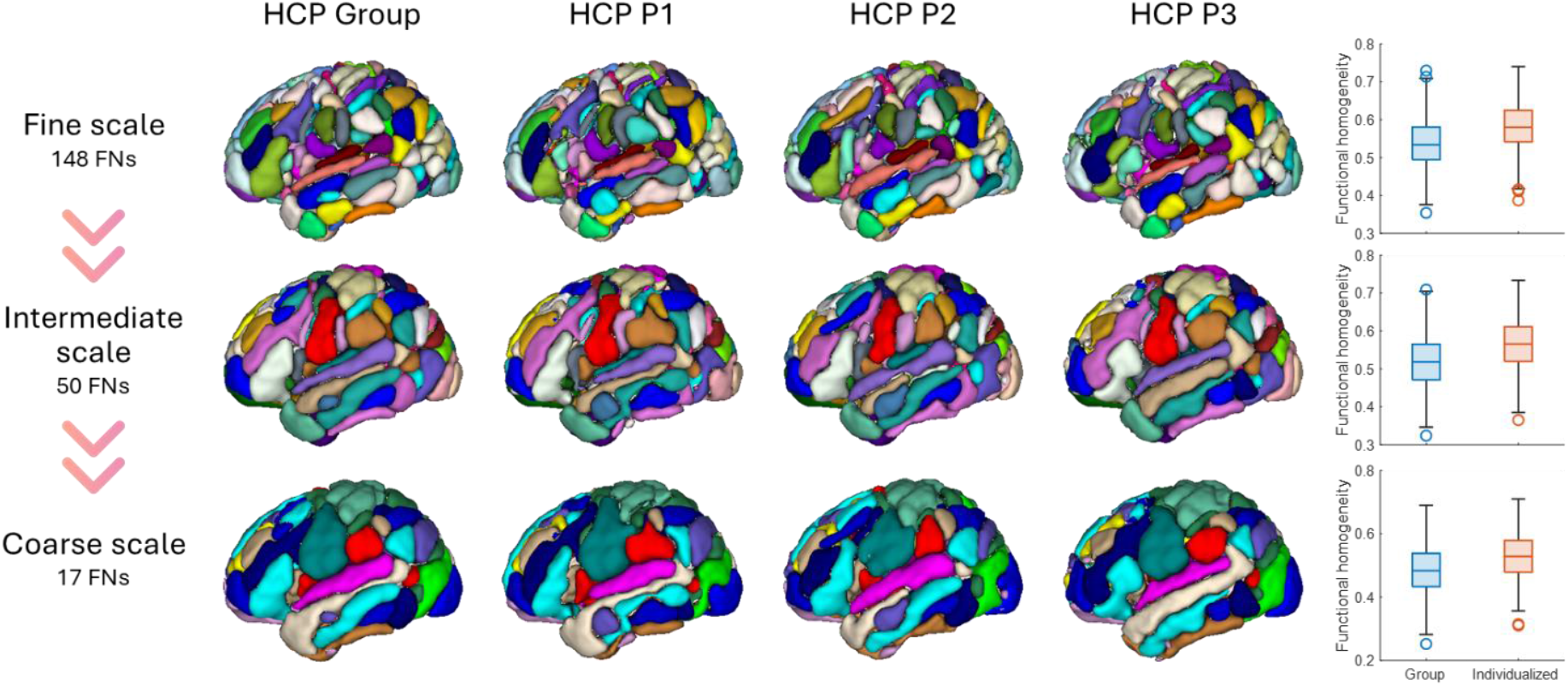
Functional parcellations derived from the group average FNs, along with those of three randomly selected HCP testing individuals (P1, P2, and P3) at fine, intermediate, and coarse scales with 148, 50, and 17 FNs, respectively. The functional parcellation was computed as a winner-take-all mapping from the FN coefficients, with colors denoting different FNs. It is worth noting that no color correspondence exists across different scales. Individualized FNs had significantly higher within-network functional homogeneity compared with the group level FNs (*p* < 0.05, Wilcoxon signed rank test).

The FNs also demonstrated clear inter-individual variation in their location, shape, and size across all the scales. Specifically, the FNs located in the association cortex exhibited substantial variability across individuals, including the dorsal attention, frontoparietal, default mode, and ventral attention networks. In contrast, FNs in the somatomotor, limbic, and visual networks showed less variability. This pattern was consistent across all scales (Fig. S4).

The functional homogeneity measures of individualized FNs were significantly higher than those of their group-level counterparts at all scales (*p* < 0.05, Wilcoxon signed rank test), indicating that individualized FNs identified by the DL model better capture the individual-specific functional organization at different spatial granularities along the hierarchy (Fig. 2 right).

### Inter-individual variability in the hierarchical structure of FNs

We investigated the hierarchical structure of FNs across scales in terms of the individualized HCs, in addition to examining the spatial topography of individualized multi-scale FNs (Fig. 2 and Fig. S4). The overall hierarchical structure of FNs across scales revealed a nested structure (Fig. 3a). This structure characterizes the segregation and integration of FNs, as quantified by the HCs computed by the DL model (Fig. 1a and 1d). Notably, most inter-scale connections occurred between FNs within the same functional module, as defined by alignment with the Yeo 7-network atlas [45]. Some finer-scale FNs were involved in multiple coarse-scale FNs in the hierarchy. For example, several ventral attention FNs at the fine scale linked to both ventral attention and somatomotor networks at the intermediate scale (Fig. 3a and Fig. 3b left).

**Fig. 3.**
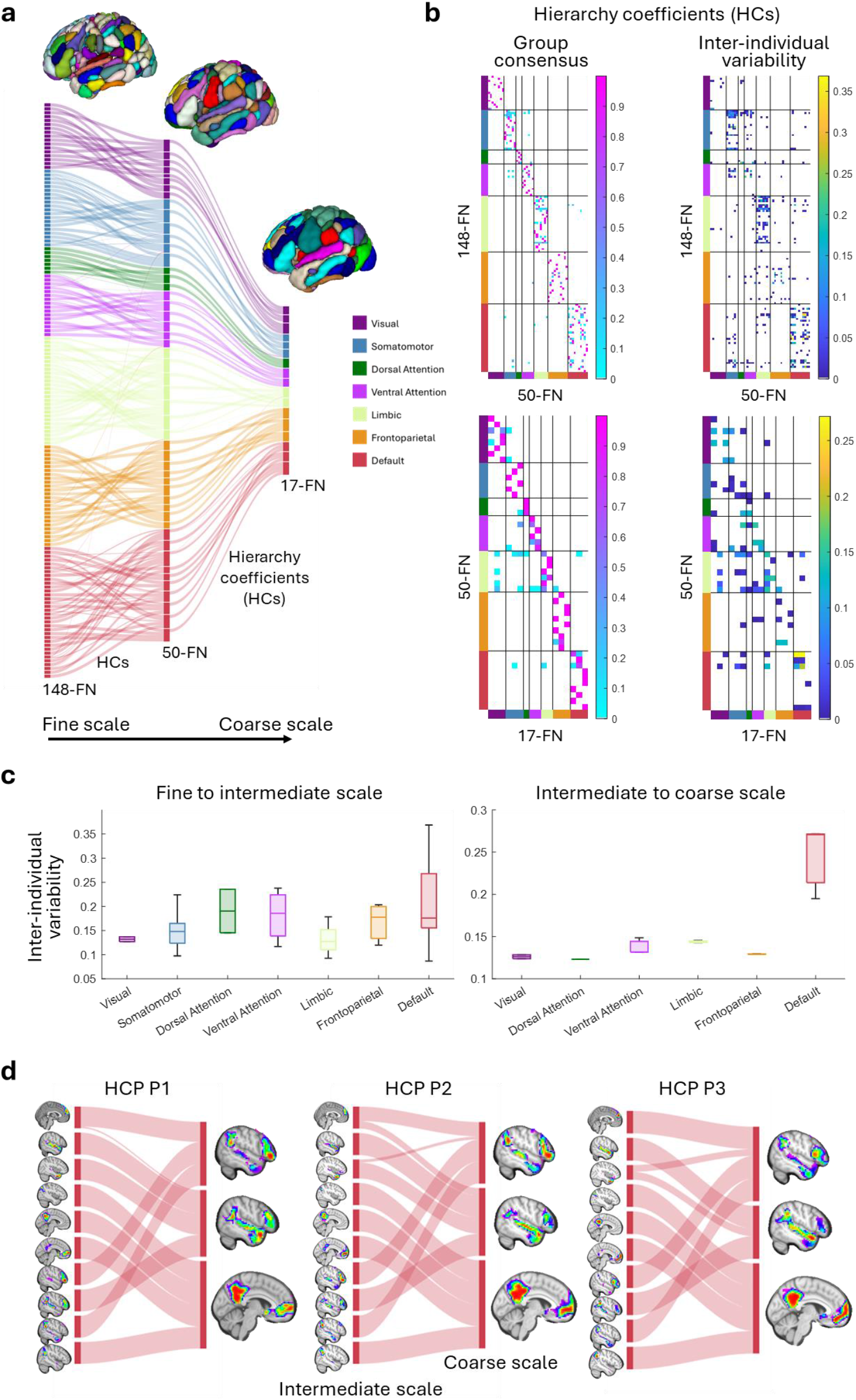
Hierarchical organization of FNs across three scales. (**a**) Group-consensus hierarchical structure illustrates the aggregation of fine-grained FNs into coarser scales, annotated and ordered according to the Yeo 7-network atlas. Each FN is assigned to the functional module with the greatest spatial overlap. (**b**) Group-consensus HCs (average HCs across individuals) and the inter-individual variability in HCs (standard deviation of HCs across individuals). (**c**) The distribution of inter-individual variability in HCs across functional modules. (**d**) Inter-individual variability is illustrated for both the FN topography (warmer colors representing higher FN loadings) and hierarchy (link width indicating HC magnitude) of the default mode network at the intermediate and coarse scales of three randomly selected individuals.

Examination of the individualized HCs revealed prominent variability in HCs between two consecutive scales across individuals (Fig. 3b right). As the HCs were mainly between FNs within the same functional module across scales, we further summarized the distribution of inter-individual variability at the functional module level (Fig. 3c). Specifically, greater inter-individual variability in HCs was observed within the default mode, dorsal attention, ventral attention, and frontoparietal networks for the fine-to-intermediate scale aggregation, while variability was most prominent in the default mode networks for the intermediate-to-coarse scale aggregation. These findings suggest that the inter-individual variation in the hierarchical structure of FNs is more prominent in the heteromodal association regions than in the unimodal regions across all scales. Furthermore, the HCs exhibited greater inter-individual variability at the fine end of the hierarchy (fine-to-intermediate) compared to the coarse end (intermediate-to-coarse).

Given the high variability in the hierarchical structure of default mode networks, we visualized the spatial topography of FNs within the default mode at the intermediate and coarse scales and their individual hierarchical structures (Fig. 3d). We observed distinct variations in spatial topography, particularly in the medial view of the coarse-scale FNs. Additionally, there were notable inter-individual differences in the interconnection pattens and magnitudes of hierarchical connections, especially between the lateral components. These findings highlight the DL model’s ability to capture the inter-individual differences in the multi-scale organization of FNs, including both the spatial topography of FNs across scales and their hierarchical structures.

### Individualized hierarchy coefficients of FNs capture sex differences

Having established the inter-individual variability in the hierarchical structure of FNs, we next investigated whether these individual variations relate to biological phenotypes at the individual level. We first examined whether individualized HCs encode sex differences in a classification setting for distinguishing males from females through multivariate pattern analysis. Specifically, we trained linear support vector machine (SVM) classifiers for distinguishing males from females under a two-fold cross-validation setting, with individualized HCs as features. The SVM models were able to classify unseen individuals as male or female with average AUCs (area under the receiver operating characteristic curve) of 0.679, 0.641, 0.705 when using fine-to-intermediate, intermediate-to-coarse, and multi-scale HCs as features respectively, demonstrating significant sex differences in HCs across individuals (Fig. 4a, all *p* < 0.001, permutation test). The multi-scale HCs were most informative in capturing the sex differences (Fig. 4a, ‘HCs_all_’), particularly when compared with the HCs between intermediate and coarse scales (‘HCs_50→17_’, *p* = 0.035, corrected *t*-test [48, 49]). No statistically significant differences in classification performance were observed between models ‘HCs_148→50_’ and ‘HCs_50→17_’.

**Fig. 4.**
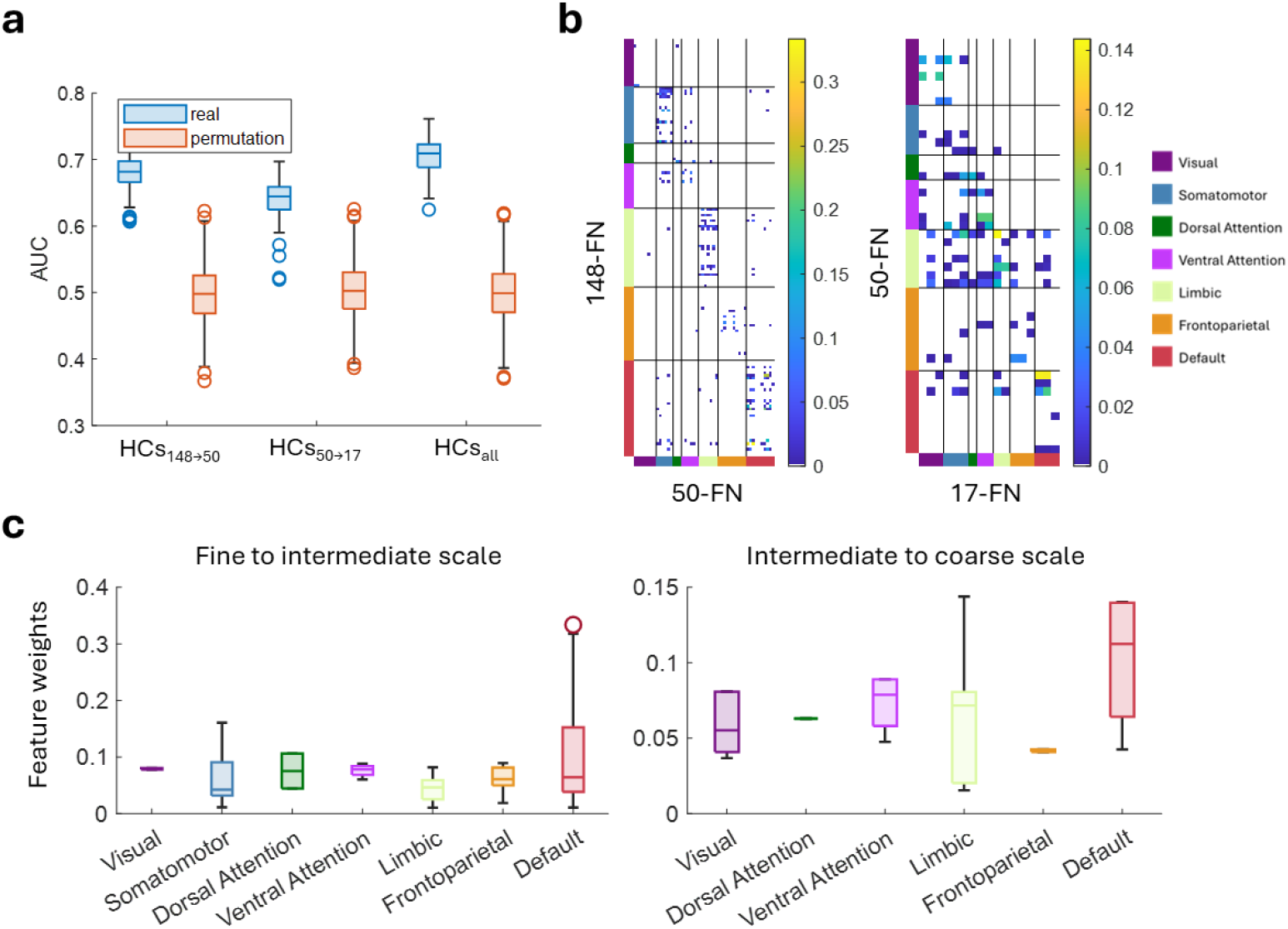
Sex differences in the hierarchical structure of FNs. (**a**) Classification performance of SVM models to classify individuals as male or female using inter-scale (‘HCs_148→50_’: fine-to-intermediate, ‘HCs_50→17_’: intermediate-to-coarse) and multi-scale (‘HCs_all_’) HCs as features. Box plots show the distribution of AUC (area under the ROC curve) measures of 100 repetitions of two-fold cross-validation. (**b**) Feature importance (the absolute Haufe-transformed feature weights [47]) of all HCs in the multi-scale HCs based classification models (‘HCs_all_’). (**c**) The distribution of feature importance of multi-scale HCs (‘HCs_all_’) across functional modules, revealing that modules including the default mode, dorsal attention, and ventral attention contributed the most to the multivariate model predicting individual sex.

To elucidate which functional modules contributed the most to the classification based on HCs, we examined the feature importance of all HCs based on the classification models and its distribution across functional modules (Fig. 4b and 4c). This analysis revealed that variability in the hierarchical structures of the default mode and dorsal attention networks (fine-to-intermediate scale) and the default mode and ventral attention networks (intermediate-to-coarse scale) contributed most to sex classification, indicating their greater importance for distinguishing males from females.

### Individualized hierarchy coefficients of FNs are associated with brain development

We also evaluated the generalizability of the DL model to individuals whose fMRI scans were collected with acquisition parameters different from the training data. We applied the model built on the HCP training cohort to fMRI data from the Philadelphia Neurodevelopmental Cohort (PNC) [50], consisting of 969 individuals (ages from 8 to 23 years). The DL model successfully identified the multi-scale hierarchical FNs for each individual and captured the inter-individual differences in spatial topography of FNs at all scales and the hierarchical structure of FNs between scales (Fig. S5), indicating good generalization.

To assess the biological relevance of the inter-individual variations in HCs identified by the DL model in the external PNC dataset, we investigated whether individualized HCs could capture brain developmental changes in the hierarchical organization of FNs. We utilized ridge regression with individualized HCs as features to construct predictive models for age under a two-fold cross-validation setting. The regression models achieved promising prediction performance when applied to unseen individuals with average correlation coefficients of 0.209, 0.129, and 0.218 between the predicted and chronological age when using fine-to-intermediate, intermediate-to-coarse, and multi-scale HCs, respectively. These results demonstrated significant developmental changes in HCs across individuals (Fig. 5a, all *p* < 0.001, permutation test). The multi-scale HCs were most informative for capturing these developmental changes (Fig. 5a, ‘HCs_all_’), particularly when compared with HCs between the intermediate and coarse scales (‘HCs_50→17_’, *p* = 0.007, corrected *t*-test). No statistically significant differences in prediction performance were observed between models ‘HCs_148→50_’ and ‘HCs_50→17_’. Feature importance analysis of the multi-scale HCs regression models revealed that variability in the hierarchical structures of the frontoparietal, ventral attention, and dorsal attention networks (fine-to-intermediate scale) and the visual and default mode networks (intermediate-to-coarse scale) contributed most to the prediction, indicating stronger correlation with brain maturation in youth (Fig. 5b and 5c).

**Fig. 5.**
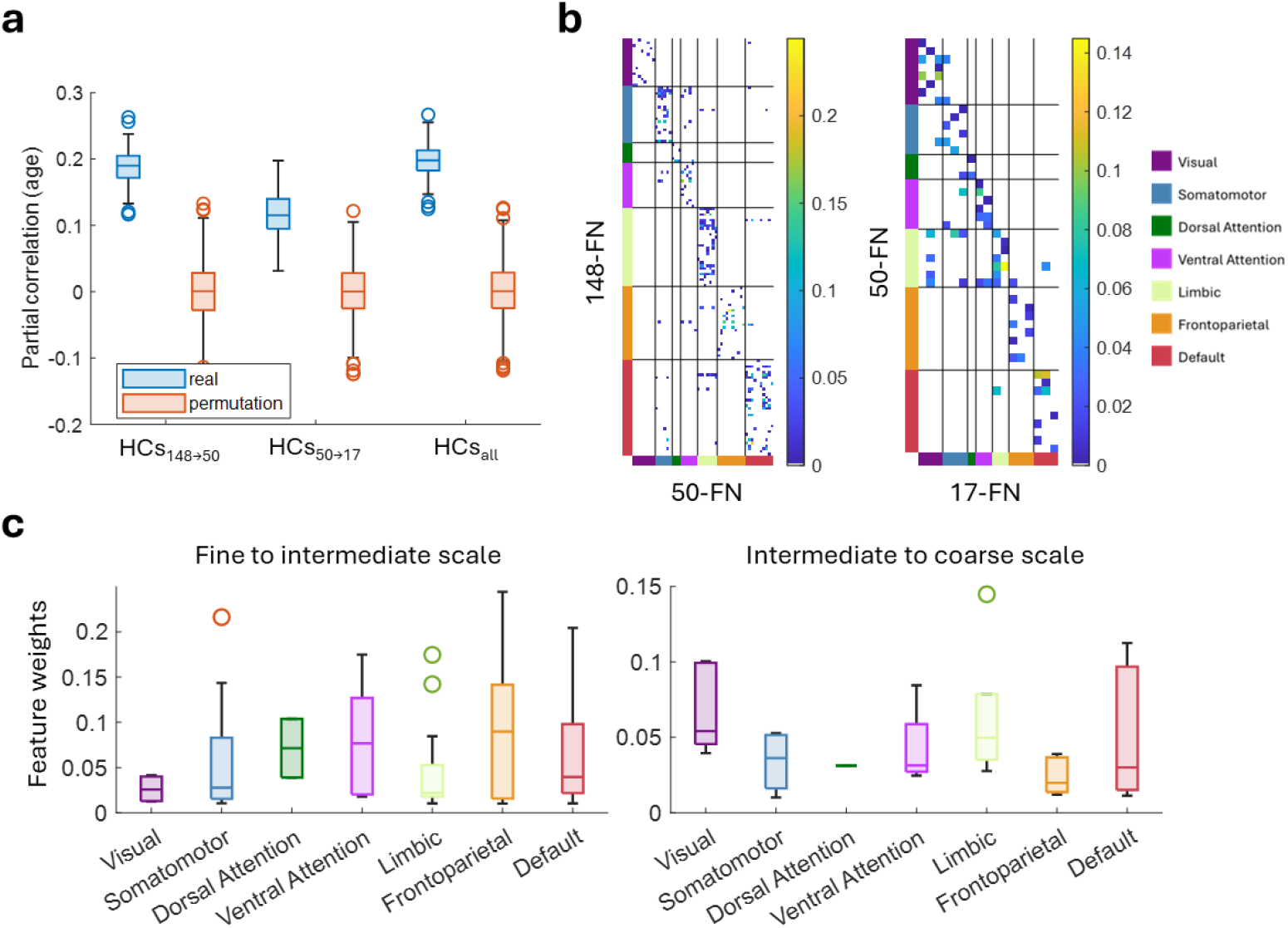
Association of the individualized hierarchy structure of FNs with the brain development in youth. (**a**) Age prediction performance using inter-scale (‘HCs_148→50_’: fine-to-intermediate scale, ‘HCs_50→17_’: intermediate-to-coarse scale) and multi-scale (‘HCs_all_’) HCs. Partial correlation coefficients between the chronological and predicted age with sex and head motion as covariates were used to measure the prediction performance. (**b**) Feature importance of all HCs in the multi-scale HCs based age prediction model (‘HCs_all_’). (**c**) The distribution of feature importance in the age prediction model (‘HCs_all_’) across functional modules.

### Individualized hierarchy coefficients of FNs distinguish healthy individuals from schizophrenia patients

Having confirmed the generalizability of the DL model trained on the HCP cohort to the external PNC dataset of healthy individuals with different age ranges, we investigated its applicability to fMRI scans of individuals with neuropsychiatric illness, aiming to identify biologically meaningful differences between healthy individuals and patients. We applied the DL model to fMRI data from an additional dataset comprising 101 healthy individuals and 94 schizophrenia (SCZ) patients. The DL model successfully identified the hierarchical multi-scale FNs for each individual and captured the inter-individual differences in spatial topography of FNs at all scales and the hierarchical structure of FNs between scales (Fig. S6).

To examine the biological relevance of individual-individual variation of HCs to schizophrenia, we investigated whether the individualized HCs of FNs were sensitive to disease-related alterations in the hierarchical structure of FNs. Using linear SVM classifiers with individualized HCs as features and two-fold cross-validation, we classified healthy individuals and SCZ patients, achieving average AUCs of 0.623, 0.650, and 0.650 using fine-to-intermediate, intermediate-to-coarse, and multi-scale HCs, respectively. These results demonstrated significant schizophrenia-related changes in HCs (Fig. 6a, all *p* < 0.02, permutation test). The multi-scale HCs and the intermediate-to-coarse scale HCs were more informative for capturing the disease-related changes (Fig. 6a, ‘HCs_all_’ and ‘HCs_50→17_’) compared with those of the fine-to-intermediate scale (‘HCs_148→50_’). Feature importance analysis of the classifiers of multi-scale HCs revealed that variability in the hierarchical structures of the dorsal attention and default mode networks (fine-to-intermediate scale) and the default mode and visual networks (intermediate-to-coarse scale) contributed most to the classification, indicating their greater discriminative power for distinguishing patients from healthy individuals (Fig. 6b and 6c).

**Fig. 6.**
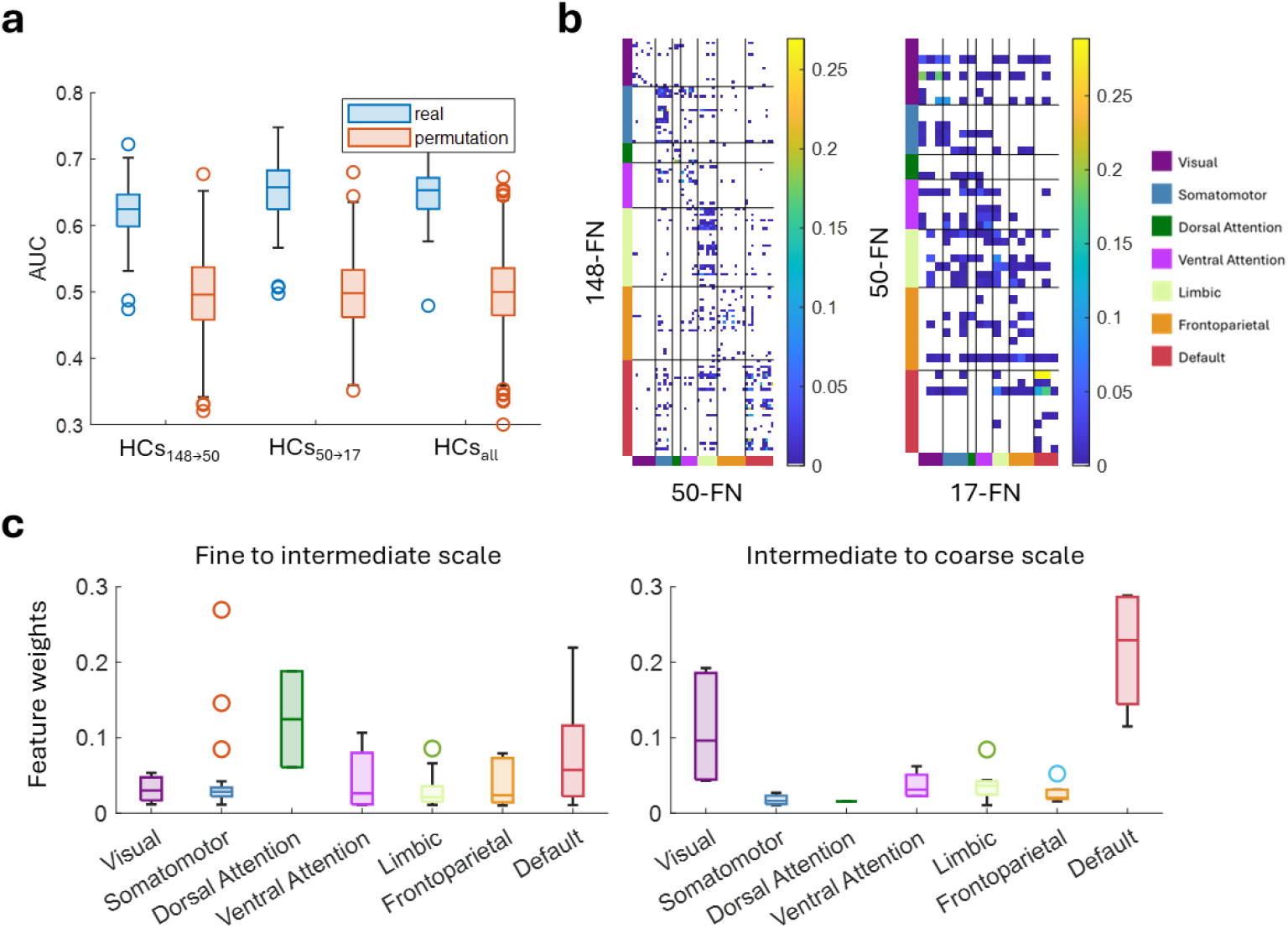
Differences in individualized hierarchy structure of FNs between healthy controls and schizophrenia (SCZ) patients. (**a**) AUCs of the classification (healthy controls vs. SCZ patients) using inter-scale (‘HCs_148→50_’: fine-to-intermediate scale, ‘HCs_50→17_’: intermediate-to-coarse scale) and multi-scale (‘HCs_all_’) HCs as features. (**b**) Feature importance of all HCs in the multi-scale HCs based classification model (‘HCs_all_’). (**c**) The distribution of feature importance in the classification model (‘HCs_all_’) across functional modules.

## Discussion

The brain is a multi-scale system with hierarchical organization that supports functional segregation and integration [28-36], exhibiting inter-individual variation in the spatial topography of FNs at different scales [1-10, 37, 51]. Characterizing these individualized multiscale FNs and their hierarchical structure is crucial for a comprehensive understanding of hierarchical functional neuroanatomy. In this study, we present a novel self-supervised DL method for end-to-end identification of individualized multiscale FNs from fMRI data. Our method learns intrinsic representations of fMRI data in a low-dimensional latent space, capturing effective embeddings of fine-grained FNs and their hierarchical structures across multiple scales, thus facilitating the computation of individualized multiscale FNs by jointly optimizing functional homogeneity of FNs at multiple scales. We demonstrate that our DL model identifies multiscale FNs with nested structure at the individual level, capturing inter-individual variation in both spatial topography and hierarchical structure of FNs. Furthermore, we demonstrate that this inter-individual variation in the hierarchical structure of FNs is associated with biological phenotypes including sex, brain development, and schizophrenia status.

Conventional approaches to characterizing multi-scale brain functional organization rely on functional parcellations or FNs computed for each scale separately [52-54], thus neglecting the hierarchical relationships between functional units across scales, despite evidence of nested structure between fine-grained and coarse FNs [37]. While clustering and module detection algorithms can identify hierarchical organization of FNs [30-34], their performance depends on pre-defined network nodes or FNs. Recursive decomposition strategies depict hierarchical structure of multi-scale FNs by dividing coarse-scale FNs into fine-grained FNs recursively [43]. However, the stepwise strategy optimizes FNs at each scale separately, neglecting the interactions of FNs between scales during the optimization procedure. Critically, both the module detection and recursive decomposition strategies capture only group-level hierarchy of FNs, failing to directly model inter-individual variation in hierarchal structure.

Unlike existing studies, our method is developed under the paradigm of predictive modeling, facilitating joint learning of the intrinsic representation of FNs and its nested structure across scales from fMRI data to construct a holistic hierarchical organization of FNs at the individual level. Our model is trained to optimize the FNs at multiple scales jointly with the learned HCs linking FNs between scales, therefore the optimization is applied to the hierarchical organization globally. Moreover, our model is trained solely based on self-supervised information, including functional homogeneity and network balance, requiring no external guidance and enabling training on any available fMRI data. After training, the model identifies individualized multi-scale FNs of unseen individuals in a single forward-pass computation without extra optimization. These computed multi-scales FNs can then be utilized for quantitative analysis of both spatial topography and inter-scale nested structure of FNs.

The proposed deep learning model identified nested multiscale FNs in unseen individuals from both internal and external testing datasets, demonstrating its generalization and broad applicability. These identified FNs were consistent with established FNs and demonstrated significant inter-individual variation in the spatial topography of FNs across all scales. Moreover, we found the inter-individual variation in the hierarchical structure of FNs, which was most predominant within functional modules of the heteromodal cortex, including the default mode, ventral attention, dorsal attention, and frontoparietal networks. Furthermore, this inter-individual variation was biologically meaningful, demonstrating significant associations with multiple phenotypes, including sex, brain development, and neuropsychiatric illness. These findings align with previous reports on inter-individual differences in FNs’ spatial topography related to sex [11] and brain development [3], suggesting that variations in topography of large-scale FNs may arise from the interplay of topography and inter-scale hierarchical structure of FNs. Multivariate pattern analyses further revealed that multi-scale hierarchy measures more closely linked with biological phenotypes than those of between-scale organizations, indicating that multi-scale measures of the hierarchical organization are more sensitive to phenotype-related differences than single-scale measures, given the difficulty in determining an optimal scale and the potential for these differences to manifest at specific scales [37].

While the individualized hierarchical FNs obtained by the DL model are promising, several open questions remain. First, the number of scales and the number of FNs at each scale were determined through a combination of data-driven techniques and prior knowledge. While the “ground-truth” hierarchical organization of functional networks remains an open question, future work should explore other configurations. Second, the quantitative analyses in the current study focused on the inter-individual variation in the hierarchy coefficients of FNs. Given the findings on inter-individual variation in FN topography [3, 11, 13, 37, 55], an integrative analysis of individual variation in both FN topography and hierarchical structure will facilitate more comprehensive understanding of the links between individualized brain functional organization and biological and clinical phenotypes. Incorporating additional measures, e.g., functional connectivity, also presents a compelling avenue for future research. Third, while the DL model demonstrated generalizability across fMRI data from different cohorts, further research is needed to investigate the impact of imaging acquisition parameters, such as number of time points of the fMRI scan and preprocessing pipelines, on the resulting FNs. Incorporating site effects of fMRI scans into the FN modeling is also an important direction. Finally, the DL model’s network architecture is currently determined empirically. Its performance can potentially be further improved using neural architecture search (NAS) techniques [56].

In conclusion, we developed a self-supervised DL method to identify individualized multi-scale functional networks with hierarchical structure. Evaluation of our method on multiple datasets has demonstrated the DL model’s ability to identify hierarchical FNs in healthy adults, children and adolescence, as well as patients with neuropsychiatric illness. Notably, the hierarchical FNs effectively captured inter-individual differences in both functional topography across scales and their hierarchical organization, exhibiting significant association with sex difference, brain development, and schizophrenia status. Moving forward, this approach provides a novel tool for investigating the holistic brain functional organization at individual level in a variety of functional neuroimaging studies.

## Methods

### Data acquisition and processing

Resting-state fMRI data from three cohorts were used in this study, including the Human Connectome Project (HCP) [44], the Philadelphia Neurodevelopmental Cohort (PNC) [50], and one cohort of healthy controls and schizophrenia (SCZ) patients [57, 58].

The HCP sample consisted of 1078 healthy individuals. Details of the scanning protocols of the fMRI data has been published previously [59]. Frame-wise displacement (FD) was adopted to measure head motion and participants with large head motion (mean FD > 0.25 mm) were excluded. The ICA-FIX denoised fMRI in volumetric space (left-to-right direction phase encoding) were used, which were spatially smoothed with a 6-mm full-width half-maximum (FWHM) kernel and downsampled at a spatial resolution of 3 × 3 × 3 mm^3^, so that the spatial resolution was consistent with that of fMRI data from the other two samples.

The PNC sample consisted of 969 young individuals (aged 8 to 23 years, 535 females). For each individual, fMRI scans were acquired using 3T Siemens Tim Trio whole-body scanner with single-shot, interleaved multi-slice, gradient-echo, echo planar imaging (GE-EPI) sequence sensitive to BOLD contrast (TR/TE=3000/32 ms, flip angle= 90^°^, FOV= 192 × 192 mm^2^, matrix=64 × 64, 46 slices, slice thickness/gap=3/0 mm, effective voxel resolution= 3 × 3 × 3 mm^3^, and 124 volumes). The fMRI data were preprocessed using an optimized procedure, including removal of the initial volumes, slice timing correction, 36-parameter confound regression, and band-pass filtering [60]. Individuals with large head motion (mean FD > 0.25 mm) were excluded.

The SCZ sample consisted of 195 individuals. All individuals underwent fMRI scanning with a 3.0-Tesla MR system (Discovery MR750, General Electric). A gradient-echo, single-short, echo planar imaging sequence was used to collect fMRI data (TR/TE=2000/45 ms, flip angle= 90^°^, FOV= 220 × 220 mm^2^, matrix= 64 × 64, 32 interleaved transverse slices, slice thickness/gap=4/0.5 mm, and 180 volumes). The fMRI scans were preprocessed using fMRIPrep [61] (version: 20.2.1). Individuals with large head motion (mean FD > 0.25 mm) were excluded. This sample comprised 101 healthy controls (age: 33.86 ± 10.77 years; 58 females) and 94 schizophrenia patients (age: 33.23 ± 8.03 years; 41 females). There was no significant difference between healthy controls and patients in age (*p*=0.647, two-sample *T*-test), gender (*p*=0.054, Chi-squared test), or head motion (*p*=0.307, two-sample *T*-test).

All the preprocessed fMRI data across samples were in the MNI 152 space and with a spatial resolution of 3 × 3 × 3 mm^3^.

### Deep learning framework for computing hierarchical FNs

Our self-supervised DL framework consists of a time-invariant representation learning module, a functional network learning module, and a functional hierarchy learning module (Fig. 1a). The representation learning module learns time-invariant feature maps from the input fMRI data, accounting for temporal misalignment of resting-state fMRI data across different scans (Fig. 1b). The functional network learning module, with an Encoder-Decoder architecture, computes individualized FNs at the fine scale from these feature maps (Fig. 1c). We hypothesized that learning an effective intrinsic representation of FNs in a low-dimensional latent space would facilitate the inference of hierarchical organization of FNs, and finer-scale FNs would aggregate into coarser-scale FNs following this hierarchy. The hierarchical organization was represented by a series of hierarchy coefficients (HCs) between two consecutive scales. The hierarchy learning module was designed to infer the inter-scale HCs from the low-dimensional latent representations of FNs extracted from the bottle-neck layer of the FN learning module (Fig. 1d).

The time-invariant representation module was implemented with the same architecture as used in our previous study [27], including a temporally separated convolution part and a temporal feature fusion part. The first part was implemented as a 3D convolutional layer with 16 filters and a stride of 1, applied to each time point of the fMRI scan. The second part was implemented as an element-wise average pooling layer, whose outputs were 16 average feature maps of all the time points of the fMRI scan. The kernel size of the convolutional layer is set to 3 × 3 × 3, and Leaky rectified linear unit (ReLU) activation and instance normalization [62] were adopted.

The functional network learning module was a Encoder-Decoder U-Net [27, 63], consisting of one convolutional layer with 16 filters, three convolutional layers with 32 filters and a stride of 2, three deconvolutional layers with 32, 32, and 16 filters and a stride of 2, and finally 2 convolutional layers with 16 filters. Leaky ReLU activation and instance normalization were used for all the convolutional and deconvolutional layers. One output convolutional layer was used to predict individualized FNs at the finest scale, and the number of output channels was *K*_1_. Softmax activation was used for the output layer so that the output was in range [0,1]. Each output channel was linearly scaled so that its maximum equals to 1. The kernel size in all layers was set to 3 × 3 × 3.

The functional hierarchy learning module consisted of *H* − 1 hierarchy projection (HP) branches to learn HCs between two consecutive scales *i* and *i* + 1, where *H* is the number of scales in the hierarchy (*H* = 3 in Fig. 1), and *i* = 1,2, …, *H* − 1. The branch *HP*_*l,l*+1_ was implemented as a subnetwork consisting of two convolution layers with *K*_*l*_ filters and a stride of 1, two fully connected (FC) layers with a latent dimension of 64, and one output FC layer with a dimension of *K*_*l*+1_, to infer the FN relationships between scales *l* and *l* + 1. The features maps from the second convolutional layer were flatten across the spatial dimension and fed into the FC layers. The latent functional embeddings from the bottleneck layer of the FN learning module were used as input to each branch *HP*_*l,l*+1_, whose output was a hierarchy coefficient matrix *HCs*_*l,l*+1_ with a size of *K*_*l*_ × *K*_*l*+1_, where *K*_*l*_ is the number of FNs at scale *l*. The entry *HCs*_*l,l*+1_(*m, n*) indicates the membership of the *m*-th FN at scale *l* to the *n*-th FN at scale *l* + 1. Leaky ReLU activation and instance normalization were used for two convolutional layers. The kernel size in convolutional layers was set to 3 × 3 × 3. Leaky ReLU activation was used for two FC layers, and Softmax activation was used for the output FC layer.

With the individualized FNs at the finest scale and hierarchy coefficient matrices between all neighboring scales, the finer-scale FNs are aggregated into coarser-scale FNs according to the learned hierarchy coefficients to construct the individualized, multi-scale FN hierarchy. The framework can be extended to more scales straightforwardly.

### Self-supervised loss function for training the DL model

A self-supervised loss function was utilized for training the DL model in an end-to-end way to optimize the data fitting performance of the FNs across all scales jointly under a matrix factorization setting, with an additional regularization on the distribution of the FNs at each scale to avoid FN collapse during model training. Since the loss function does not rely on any other external supervision information, the DL model was optimized based on input data alone, referred to as self-supervised.

Given a group of *n* individuals, each having 4D fMRI data 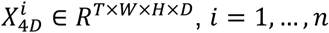, where *W, H*, and *D* are width, height, and depth of each 3D volume respectively and *T* is the number of time points, the proposed DL model takes the 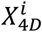 as input and identify its corresponding FNs 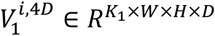 at the finest scale and inter-scale HCs 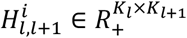 (*l* = 1,2, …, *H* − 1) as output. For calculating the loss function, the volumetric data 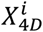 and 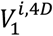 are flattened across the spatial dimension and reshaped as matrices *X*^*i*^ ∈ *R*^*T*×*S*^ and 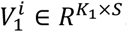, where *S* = *W* × *H* × *D* is the number of voxels within the brain. Then, the self-supervised loss function is calculated as

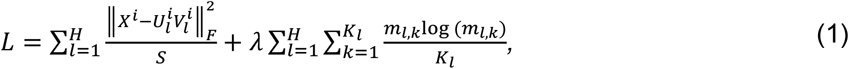

where 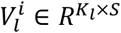 and 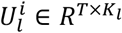 refer to *K*_*l*_ FNs and their corresponding time courses at scale 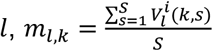 is the proportion of all voxels in the *k*-th FN at scale *l, H* is the number of scales in the hierarchy of FNs. The first term measures the data-fitting error when using FNs to approximate the original fMRI data, the second term measures the balance of FN size to avoid tiny or empty FNs, and λ is the trade-off hyperparameter. More specifically, FNs at the finest scale 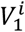 is output by the DL model, and 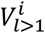 is obtained by 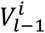 and the HCs 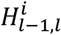 as 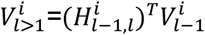, followed by FN-wise normalization as 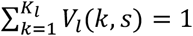, where 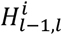 is also the output of the DL model. The time courses 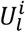 is computed as 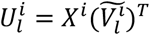, where 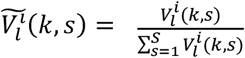.

We used fMRI data from *n* individuals of the training cohort to minimize the loss function in Eq. (1) for optimizing the network weights of the DL model. After training, the model can be applied to fMRI data of new individuals to identify their individualized hierarchical FNs in a single forward pass, and the individualized hierarchy coefficients can be used as quantitative measures to investigate the inter-individual variation in the hierarchy of brain functional organization.

### Number of FNs from fine to coarse scales

A hierarchy of FNs with three scales was identified and analyzed, consisting of 148, 50, and 17 FNs respectively from fine to coarse scales. The number of FNs at the finest scale was estimated using Laplace approximation as implemented in MELODIC from FSL [64]. Specifically, fMRI data of 100 subjects from the training cohort were randomly selected, based on which the number of FNs was estimated using the Laplace approximation technique. The procedure was repeated 50 times, and the median of the estimated numbers of FNs was selected. The final number of FNs at the finest scale was set to 148. The number of FNs at the coarsest scale was set to 17, as widely used in the literature [45]. The number of FNs at the intermediate scale was set to 50, so that the granularity of FNs decreased by a factor of 3 approximately from fine to coarse scales.

### Implementation details

Our model was implemented using Pytorch [65]. Adam optimizer was adopted to optimize the network, the learning rate was set to 1 × 10^−4^, the batch size was set to 1, and the number of epochs was set to 100 during training. One NVIDIA A40 GPU with 48 GB memory was used for training and testing. We set λ = 50 empirically in our experiments.

### Quantitative evaluation of FNs

The homogeneity measure for each FN was calculated as the weighted sum of the correlation coefficients between the time courses of all the voxels within the FN and its centroid time course, which was a weighted mean time course within the FN and the FN’s voxelwise loadings were used as weights. The median value of FN-wise homogeneity measures was used to gauge the overall functional homogeneity of FNs for each individual.

The inter-individual variability in FN topography was calculated as the averaged median absolute deviation maps of FN loadings across individuals over all FNs, as previously used in [3]. The inter-individual variability map was calculated for each scale separately.

### Multivariate predictive modeling

The HCs between two consecutive scales (‘HCs_148→50_’: 148-FN to 50-FN, ‘HCs_50→17_’: 50-FN to 17-FN) and across all scales (‘HCs_all_’: combinations of ‘HCs_148→50_’ and ‘HCs_50→17_’) were utilized as feature vectors for each individual to build inter-scale HCs and multi-scale HCs based predictive modeling. Linear support vector machine (SVM) [66] was utilized to build binary classifiers under the classification scenarios (sex classification and schizophrenia classification), and ridge regression [67] was utilized to build models for age prediction. All predictive modeling were conducted with 2-fold cross-validation, and regularization parameters in the model were optimized using nested 2-fold cross-validation within the training cohort. The regularization parameter in the predictive models was optimized in the range [2^−10^, 2^−9^, …, 2^4^, 2^5^]. The classification performance was evaluated by area under the receiver operating characteristic (ROC) curve (AUC), and the regression performance was evaluated by partial correlation coefficient between the predicted and real measures with sex and in-scanner head motion as co-variates. The prediction for each phenotype of interests was repeated 100 times. Permutation testing with 1000 permutations was utilized to test the significance of prediction performance over chance.

To elucidate the importance of different features for the prediction, we summarized the feature weights in prediction models for each phenotype. Specifically, we averaged the absolute Haufe-transformed feature weights [47] of prediction models across 100 repetitions as the feature importance and summarize their distribution across different functional modules as in the Yeo 7-network atlas.

### Statistical tests

Spin-based spatial permutation procedure [46] was leveraged to test the alignment of our functional parcellation derived from the group average FNs and the Yeo 17-network atlas. Specifically, our functional parcellation at the MNI 152 was first projected to the fsaverage space using neuromaps [68], and its parcel labels were matched to Yeo atlas labels using the Hungarian method [69]. Dice coefficient was used to measure the similarity for each pair of matched parcels, and the averaged Dice coefficient across parcels served as the overall similarity. The null distribution of the similarity measures was generated by computing the Dice coefficient between our functional parcellation and the spin rotated Yeo atlas over 1000 permutations. The true similarity measure was finally compared to the null distribution to determine the significance.

To test whether the individualized FNs had higher functional homogeneity than the group average FNs, the overall functional homogeneity of FNs was calculated for each individual using its individualized FNs and the group average FNs respectively, Wilcoxon signed rank test was then used to compare the homogeneity measures across individuals.

To test whether a model’s classification/prediction performance is better than chance, permutation tests by shuffling class label/target measure across individuals and performing the cross-validation procedure. We used a corrected *t*-test [48, 49] to compare the classification/prediction performance of two models on the same task. This test accounts for the dependency of accuracy measures from testing folds in repeated cross-validation.

## Data and code availability

The HCP data is publicly available at https://db.humanconnectome.org. The PNC data is accessible from the Database of Genotypes and Phenotypes (phs000607.v3.p2) at https://www.ncbi.nlm.nih.gov/projects/gap/cgi-bin/study.cgi?study_id=phs000607.v3.p2. The in-house cohort of healthy controls and SCZ patients will not be publicly available. Codes used for model training and testing will be available at: https://github.com/MLDataAnalytics.

## Acknowledgements

This study was supported by grants from the National Institute of Health: R01EB022573, R01AG066650, U24NS130411.

