## Supplemental materials for "Mapping individualized multi-scale hierarchical brain functional networks from fMRI by self-supervised deep learning"

[1]Center for AI and Data Science for Integrated Diagnostics (AI2D), University of Pennsylvania, Philadelphia, PA 19104, USA
[2]Center for Biomedical Image Computing and Analytics (CBICA), University of Pennsylvania, Philadelphia, PA 19104, USA
[3]Department of Radiology, University of Pennsylvania, Philadelphia, PA 19104, USA
[4]Laboratory of Psychiatric-Neuroimaging-Genetic and Co-morbidity (PNGC_Lab), Tianjin Anding Hospital, Tianjin Mental Health Center of Tianjin Medical University, Tianjin, 300222, China
[5]Chinese Institute for Brain Research, Beijing, 102206, China
[6]Department of Psychiatry, University of Pennsylvania, Philadelphia, PA 19104, USA
[7]Penn/CHOP Lifespan Brain Institute, University of Pennsylvania, Philadelphia, PA 19104, USA
[8]Department of Neurology, University of Pennsylvania, Philadelphia, PA 19104, USA
[9]Penn Statistics in Imaging and Visualization Center, University of Pennsylvania, Philadelphia, PA 19104, USA
[10]Department of Biostatistics, Epidemiology and Informatics, University of Pennsylvania, Philadelphia, PA 19104, USA
[11]Center for Brain Imaging and Stimulation, University of Pennsylvania, Philadelphia, PA 19104, USA
[12]Penn Brain Science, Translation, Innovation, and Modulation Center, University of Pennsylvania, Philadelphia, PA 19104, USA
[13]Center for Neuromodulation in Depression and Stress, University of Pennsylvania, Philadelphia, PA 19104, USA
[14]Departments of Neuroscience, Neurosurgery and Bioengineering, University of Pennsylvania, Philadelphia, PA 19104, USA
[15]Penn Lifespan Informatics and Neuroimaging Center, University of Pennsylvania, Philadelphia, PA 19104, USA

### Supplemental materials

Supplemental figures Fig. S1 to Fig. S6.

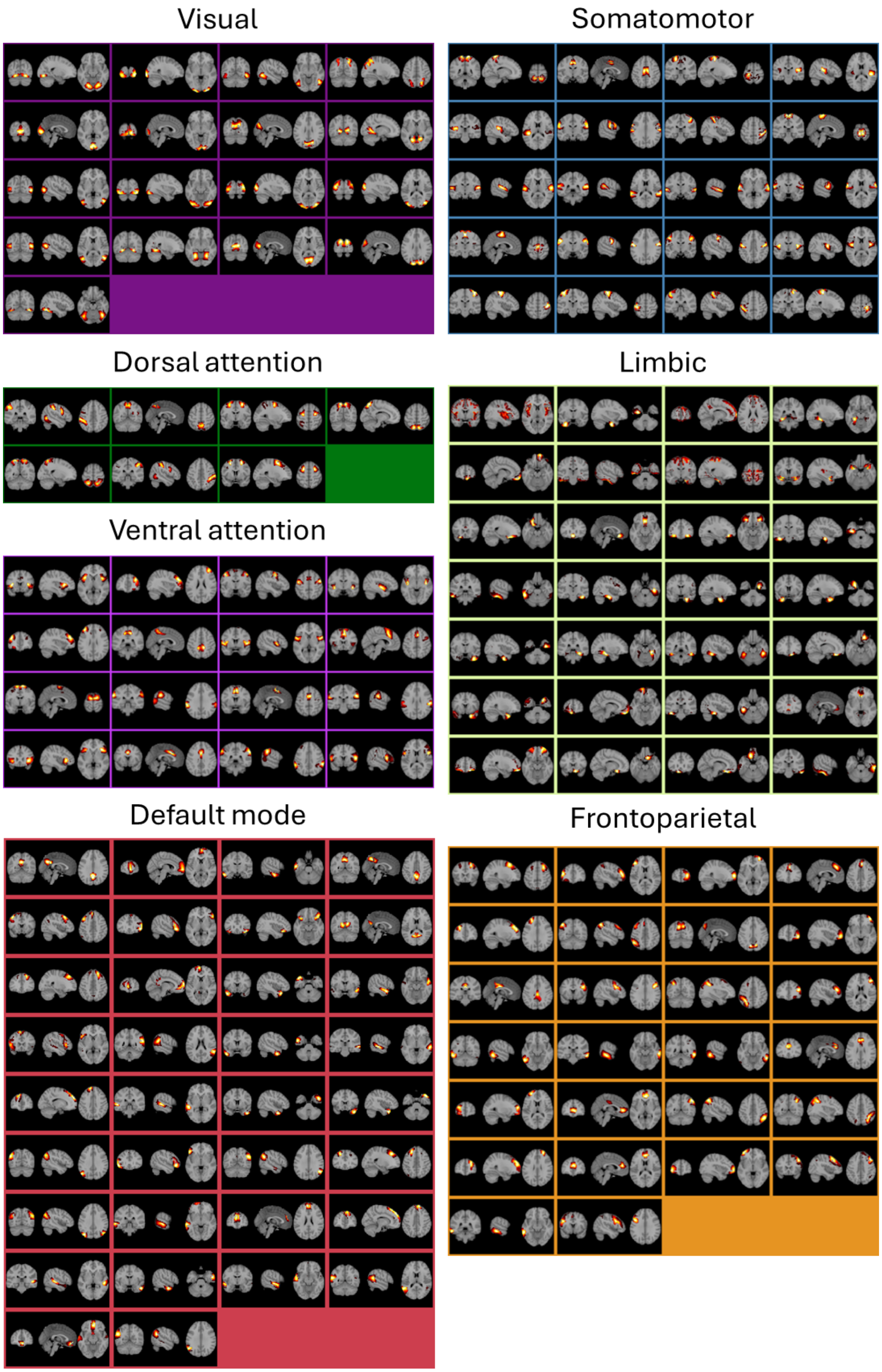


**Fig. S1**. Group-average FNs of the HCP testing individuals at the fine scale (148 FNs). FNs were annotated according to the Yeo 7-network atlas. Each FN was assigned to the functional module with the greatest spatial overlap.

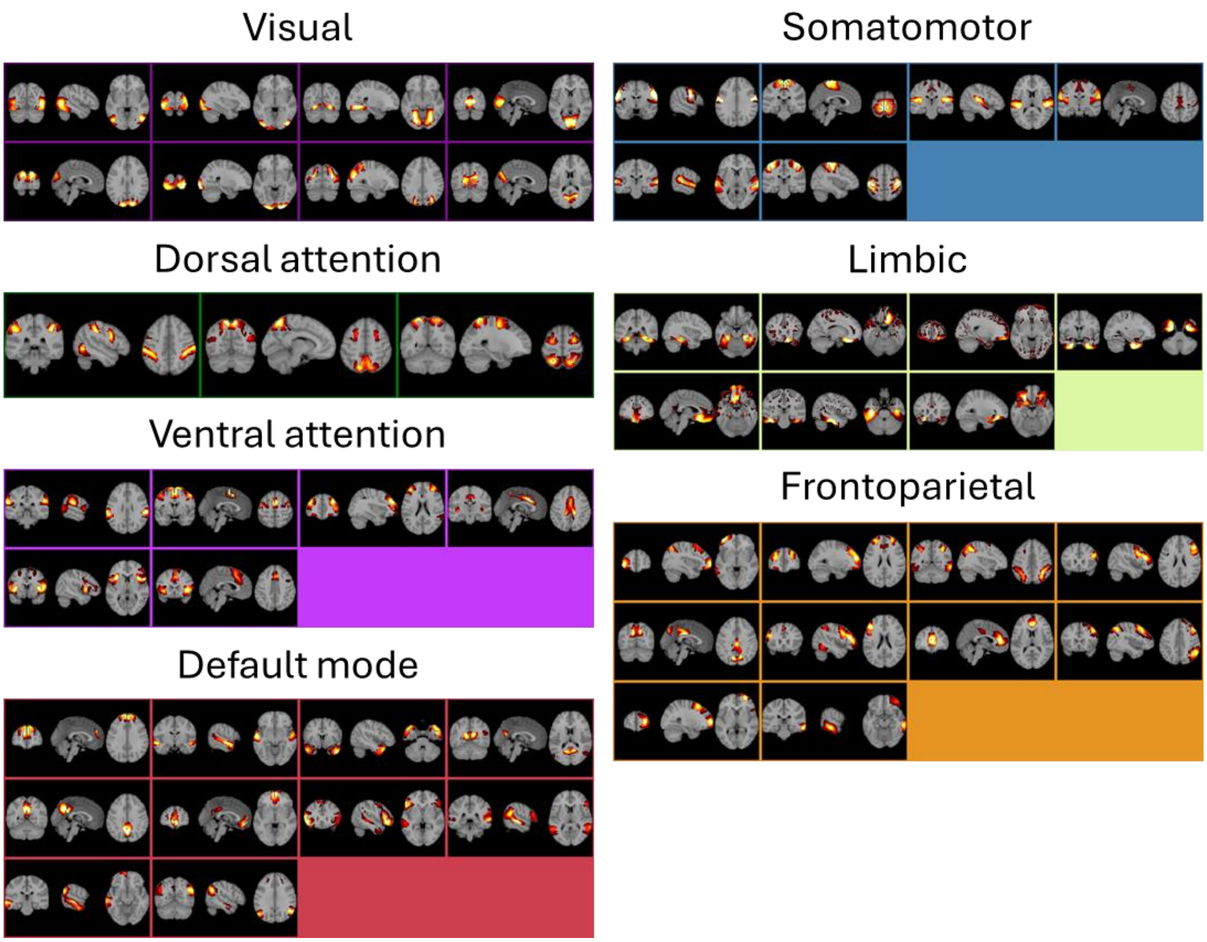


**Fig. S2**. Group-average FNs of the HCP testing individuals at the intermediate scale (50 FNs). FNs were annotated according to the Yeo 7-network atlas. Each FN was assigned to the functional module with the greatest spatial overlap.

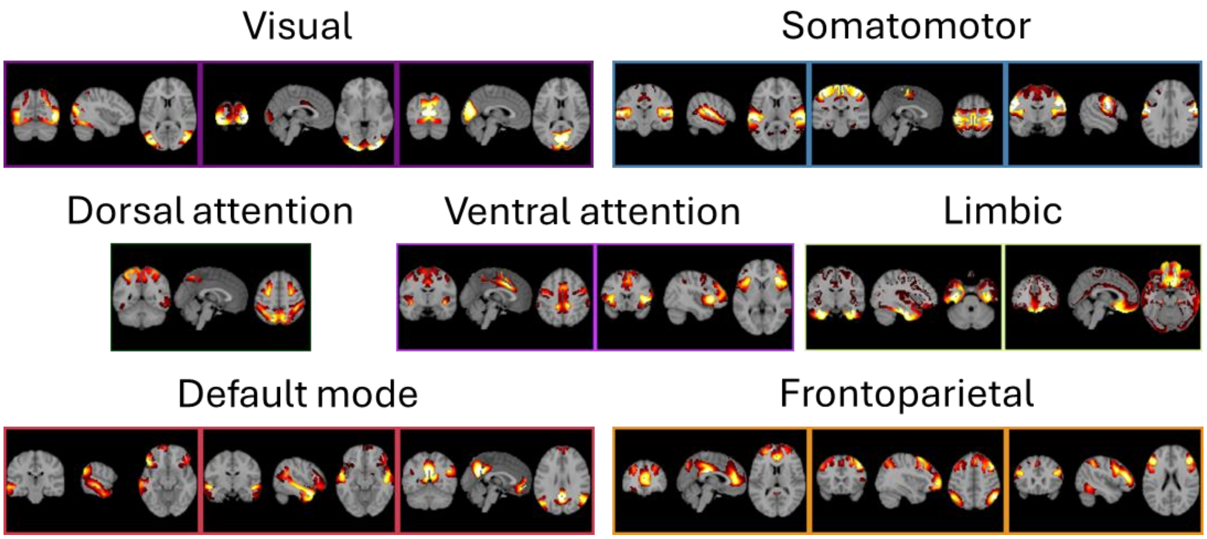


**Fig. S3**. Group-average FNs of the HCP testing individuals at the coarse scale (17 FNs). FNs were annotated according to the Yeo 7-network atlas. Each FN was assigned to the functional module with the greatest spatial overlap.

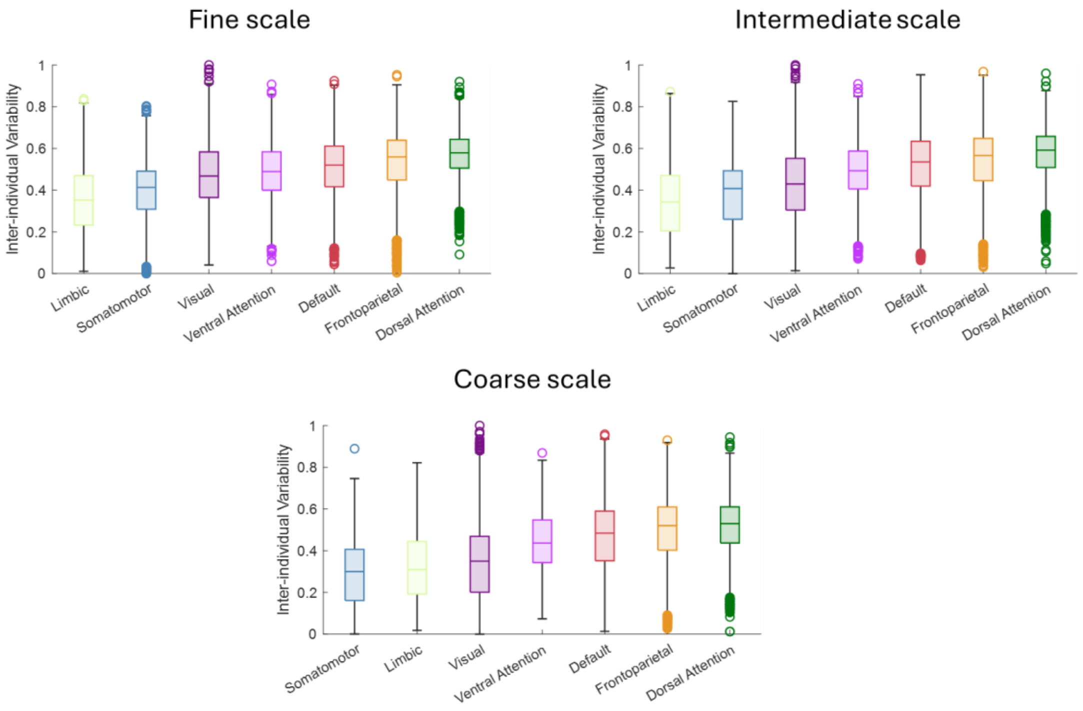


**Fig. S4**. Inter-individual variability of FN topography across functional modules at different scales. Voxel-wise variability of FN topography across individuals was summarized using the Yeo 7-network atlas. The voxel-wise variability was scaled to a range between 0 and 1 by min-max normalization for each scale.

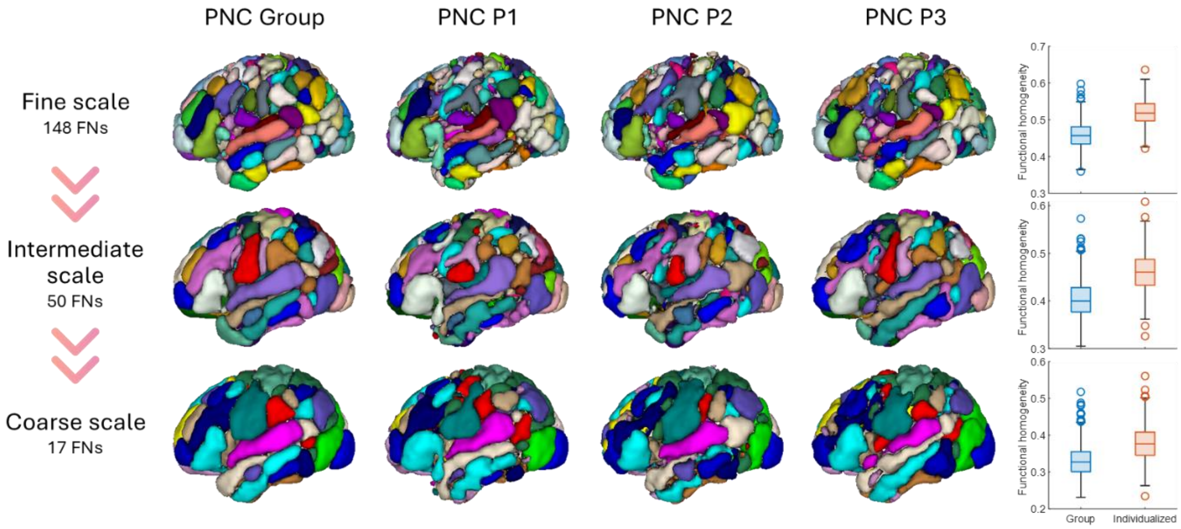


**Fig. S5.** Functional parcellations derived from group-average FNs of the PNC sample, along with those of three randomly selected PNC testing individuals (P1 to P3) at fine, intermediate, and coarse scales with 148, 50, and 17 FNs, respectively. The functional parcellation was computed as a winner-take-all mapping from the FN coefficients, with colors denoting different FNs. It is worth noting that no color correspondence exists across different scales. Individualized FNs had significantly higher within-network functional homogeneity compared with the group level FNs ($p < 0.05$, Wilcoxon signed rank test).

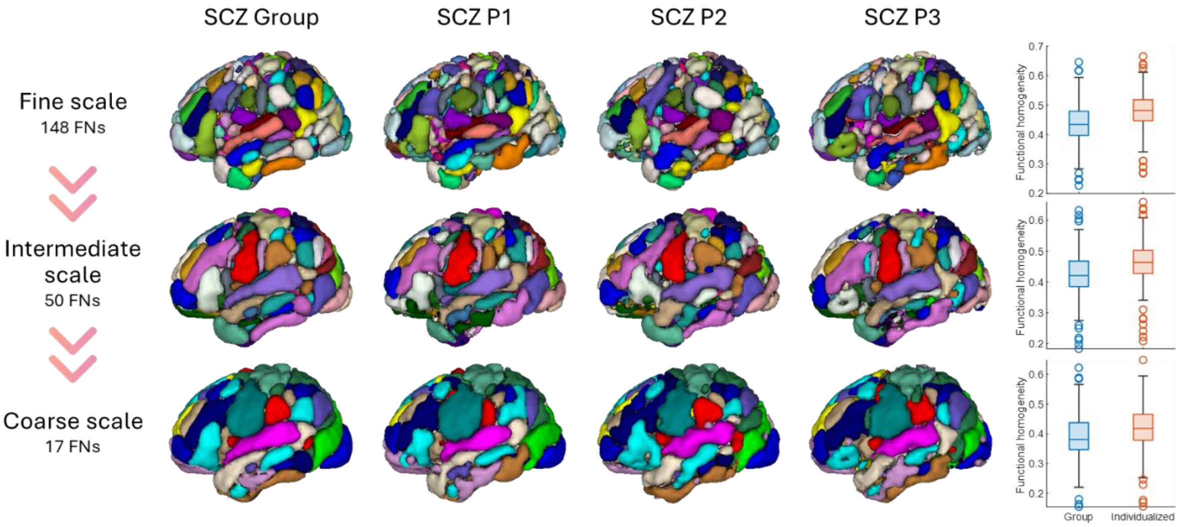


**Fig. S6.** Functional parcellations derived from group-average FNs of the SCZ sample, along with those of three randomly selected SCZ testing individuals (P1 to P3) at fine, intermediate, and coarse scales with 148, 50, and 17 FNs, respectively. The functional parcellation was computed as a winner-take-all mapping from the FN coefficients, with colors denoting different FNs. It is worth noting that no color correspondence exists across different scales. Individualized FNs had significantly higher within-network functional homogeneity compared with the group level FNs ($p < 0.05$, Wilcoxon signed rank test).
